# KCH kinesin drives nuclear transport and cytoskeletal coalescence for tip cell growth

**DOI:** 10.1101/308775

**Authors:** Moé Yamada, Gohta Goshima

## Abstract

Long-distance transport along microtubules (MTs) is critical for intracellular organisation. In animals, antagonistic motor proteins kinesin (plus end-directed) and dynein (minus end-directed) drive cargo transport. In land plants, however, the identity of motors responsible for transport is poorly understood, as genes encoding cytoplasmic dynein are missing. How other functions of dynein are brought about in plants also remains unknown. Here, we show that a subclass of the kinesin-14 family, KCH—which can also bind actin—drives MT minus end-directed nuclear transport in the moss *Physcomitrella patens*. When all four KCH genes were deleted, the nucleus was not maintained in the cell centre but was translocated to the apical end of protonemal cells. In the knockout (KO) line, apical cell tip growth was also severely suppressed. KCH was localised on MTs, including at the MT focal point near the tip where MT plus ends coalesced with actin filaments. MT focus was not persistent in *KCH* KO lines, whereas actin destabilisation also disrupted the focus despite KCH remaining on unfocused MTs. Functions of nuclear transport and tip growth were distinct, as a truncated KCH construct restored nuclear transport activity but not tip growth retardation of the KO line. Thus, our study identified KCH as a long-distance retrograde transporter as well as a cytoskeletal crosslinker, reminiscent of the versatile animal dynein.

## INTRODUCTION

Intracellular transport is a critical cellular mechanism for cell organisation in eukaryotic cells. Many cellular components, including organelles, proteins, and RNA, are transported to their appropriate positions where they specifically function in response to internal and external signals. Although it had been believed that plants predominantly utilize actin and myosin to move cellular components, recent studies have uncovered the prevalence of microtubule (MT)-dependent transport as well (Kong et al., 2015; Miki et al., 2015; Nakaoka et al., 2015; Zhu et al., 2015; Yamada et al., 2017). However, a unique feature of plant motor systems is that the genes encoding cytoplasmic dynein, the sole MT minus end-directed transporter in animals, have been lost during plant evolution. Moreover, dynein function is not limited to cargo transport, as a variety of fundamental cellular processes requires dynein, such as MT-based force generation at the cortex (Grill and Hyman, 2005; Gonczy, 2008; McNally, 2013), MT-MT crosslinking (Ferenz et al., 2009; Tanenbaum et al., 2013), and MT-actin crosslinking (Grabham et al., 2007; Perlson et al., 2013; Coles and Bradke, 2015). However, how plants execute these functions without dynein remains unanswered.

The moss *Physcomitrella patens* is an emerging model plant of cell and developmental biology, in part due to the applicability of homologous recombination and high-resolution live imaging (Cove, 2005; Cove et al., 2006; Vidali and Bezanilla, 2012). The protonemal apical cell of *P. patens* is an excellent system to study MT-based transport. MTs are predominantly aligned along the cell longitudinal axis with a characteristic overall polarity depending on cell cycle stage (illustrated in Figure 2A). Nucleus, chloroplasts, and newly formed MTs have been identified as cargo that is transported on MT tracks in protonemal cells, wherein nuclear movement was shown to be independent of actin (Miki et al., 2015; Nakaoka et al., 2015; Yamada et al., 2017). Using this model system, kinesin-ARK (armadillo repeat-containing kinesin) was first identified as a plus end-directed nuclear transporter; upon RNAi knockdown of this plant-specific, plus end-directed motor protein, the nucleus migrated towards the cell centre after cell division as normal but then moved back to the cell plate, i.e. the nucleus showed an abnormal minus end-directed motility (Figure 2A; Miki et al., 2015). It was also revealed that the non-processive, minus end-directed KCBP (kinesin-like calmodulin-binding protein)—a member of the kinesin-14 protein family—is required for minus end-directed nuclear transport. In the absence of KCBP, the nucleus could not move to the cell centre immediately after cell division, i.e. minus end-directed motility was inhibited (Figure 2A; Yamada et al., 2017). Although a single dimeric KCBP cannot take multiple steps along the MT (non-processive), clustered motors exhibit processive motility *in vitro* and *in vivo*; thus, multiple KCBP motors associated with the nuclear surface can transport the nucleus towards MT minus ends (Jonsson et al., 2015; Yamada et al., 2017). However, nuclear transport function of KCBP is limited during the latest stage of cell division, as KCBP is no longer necessary for maintaining central positioning of the nucleus during interphase. It is plausible that an additional minus end-directed motor protein that antagonises kinesin-ARK and possibly other plus end-directed kinesins is expressed in moss cells.

Minus end-directed kinesin-14 is duplicated uniquely in the land plant lineage and constitutes six subfamilies; in *Arabidopsis*, it is the most expanded family among the kinesin superfamily (Zhu and Dixit, 2011; Shen et al., 2012). KCBP belongs to class VI of kinesin-14 and transports not only the nucleus but also chloroplasts in moss (Yamada et al., 2017). In *Arabidopsis*, the cytoskeletal organisation of the trichome cell is defective in *kcbp* mutants, suggesting an additional function to nuclear/chloroplast transport (Tian et al., 2015). The class I kinesin-14 ATK (HSET/XCTK2/Ncd) conserved in animals, is essential for mitotic spindle coalescence, and drives minus end-directed transport of newly formed MTs along other MTs in the moss cytoplasm (Ambrose et al., 2005; Yamada et al., 2017). KAC/KCA is a class V kinesin-14 that no longer possesses MT affinity but has acquired an actin-binding region and regulates actin-dependent chloroplast photo-relocation movement and anchorage to the plasma membrane (Suetsugu et al., 2010; Suetsugu et al., 2012). Class III kinesin-14 is localised to the spindle in moss but appears to have lost MT-based motor activity (Miki et al., 2014; Jonsson et al., 2015). The class IV kinesin-14 TBK has a weak MT motor activity and localises to cortical MTs yet its cellular function is unknown (Goto and Asada, 2007; Jonsson et al., 2015). Class II kinesin-14 genes form a large clade in the plant kinesin family, where 9 out of 61 *A. thaliana* kinesin genes are classified into this clade (Figure 1A) and multiple activities and cellular functions have been reported. Kinesin14-II possesses the calponin homology (CH) domain in its amino-terminal region followed by dimerisation and motor domains (hereafter called KCH, which stands for kinesin with CH domain; Figure 1B). Adjacent to the motor domain, there exists an uncharacterised C-terminal extension in this subfamily that is not found in ATK or KCBP (Preuss et al., 2004; Frey et al., 2009; Shen et al., 2012). Mutant analyses have uncovered divergent functions of KCH, such as cell size regulation (OsKCH1; Frey et al., 2010), mitochondrial respiration (*Arabidopsis* KP1; (Yang et al., 2011)), and cell-to-cell movement of a transcription factor (*Arabidopsis* KinG; Spiegelman et al., 2018). However, the complete picture of KCH function has not been elucidated, since loss-of-function analysis using complete null mutants has not been conducted for this highly duplicated gene subfamily in flowering plants. In contrast, *P. patens* possesses only four *KCH* genes that are highly homologous (Figure 1A; red), suggesting that they redundantly exhibit basal functions of this kinesin subfamily.

**Figure 1.**
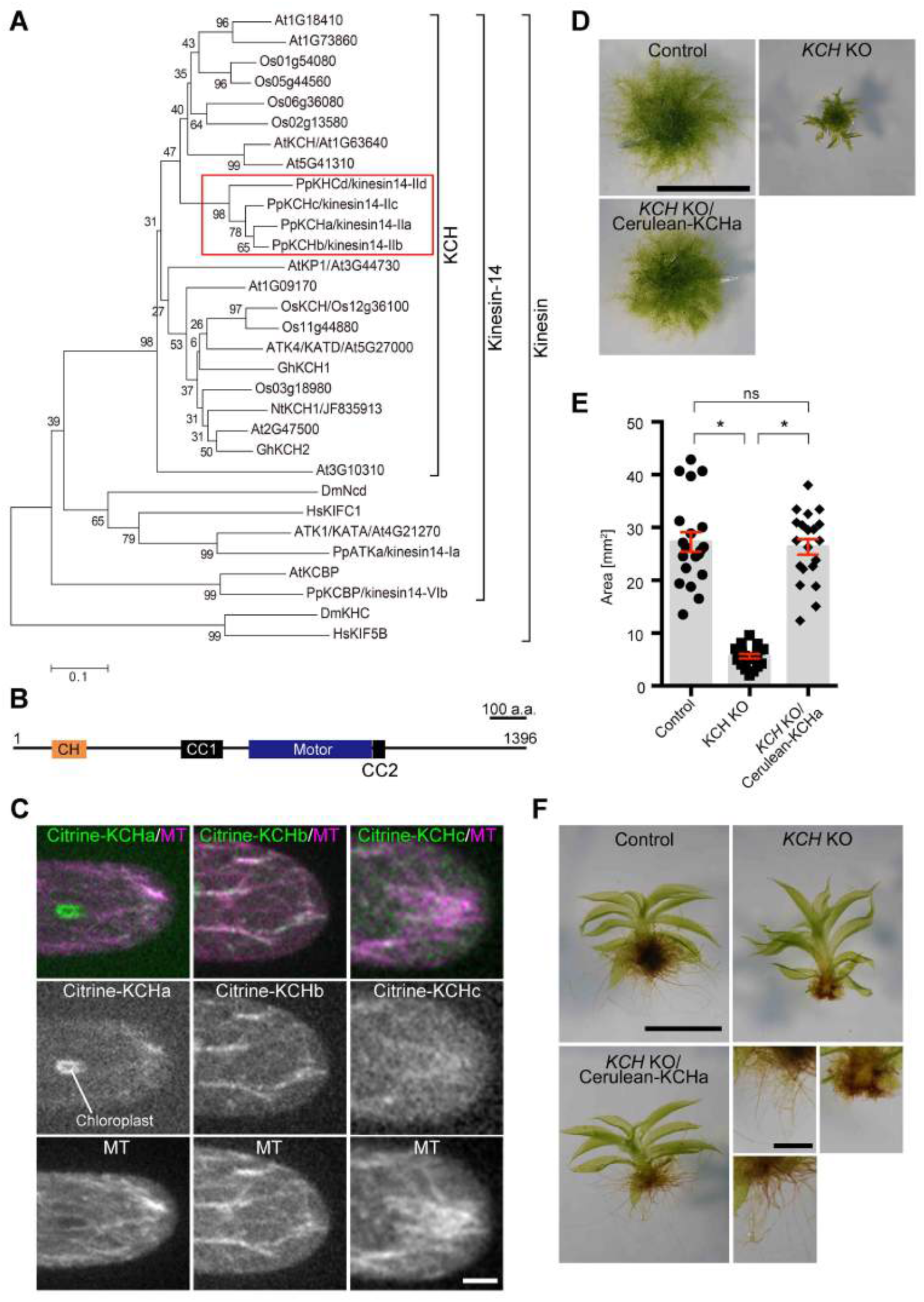
KCH is required for protonema growth, gametophore leaf morphology, and rhizoid elongation. **(A)** Phylogenetic tree of plant KCH subgroup members and other members of the kinesin family. At, *Arabidopsis thaliana*; Os, *Oryza sativa*: Pp, *Physcomitrella patens*; Gh, *Gossypium hirsutum*; Nt, *Nicotiana tabacum*; Dm, *Drosophila melanogaster*, Hs, *Homo sapiens*. Horizontal branch length is proportional to the estimated evolutionary distance. Scale bar, 0.1 amino acid substitutions per site. **(B)** Schematic diagram of domain organisation and coiled-coil (CC) prediction of P. patens KCHa (1-1396 a.a.). Calponin homology (CH) domain (106-199 a.a.) and kinesin motor domain (642-975 a.a.) were predicted by an NCBI domain search. Predicted coiled-coil with a probability > 0.7 and 21 residue windows (Coiled-Coils Prediction; PRABI Lyon Gerland) are displayed (454-567 a.a. and 978-1012 a.a.). **(C)** MT localisation of Citrine-KCHa, -KCHb, and -KCHc at the caulonemal cell tip. Image contrast was individually adjusted for each sample. Scale bar, 5 μm. **(D, E)** Colony size comparison between control (parental line expressing GFP-tubulin/histoneH2B-mRFP), *KCH* KO, and *KCH* KO/Cerulean-KCHa. Colonies were cultured for 3 weeks from the stage of protoplasts. Scale bar in (D), 5 mm. Bars and error bars in (E) represent the mean and SEM, respectively. Control, n = 20; *KCH* KO, n = 20; *KCH* KO/Cerulean-KCHa, n = 20. *P < 0.0001; ns (not significant), P > 0.7 (unpaired t-test with equal SD, two-tailed). Experiments were performed three times and the data analysed twice. The data of one experiment is displayed. (F) Gametophores and rhizoids cultured for 4 weeks on BCDAT medium. Scale bar, 1 mm (top) or 0.5 mm (bottom).

In this study, we generated a plant with a complete deletion of the *KCH* gene of *P. patens*, and provide evidence that KCH drives minus end-directed nuclear transport. Furthermore, KCH contributes to cell tip growth likely via crosslinking MTs at the apical tip. These two functions are distinct, as a KCH fragment that lacks the unusual C-terminal extension fulfils the function of nuclear transport but not tip growth. In contrast, the CH domain, which has been assumed to be the cargo (i.e. actin) binding site, was not required for either function. We propose that plant KCH is a versatile cargo transporter that also fulfils other MT-based functions, analogous to the dynein motor in animals.

## RESULTS

### Complete KCH deletion affects moss growth and morphology

The expression database (http://bar.utoronto.ca/efp_physcomitrella/cgi-bin/efpWeb.cgi) (Ortiz-Ramirez et al., 2016) and our previous localisation analysis of KCH-Citrine fusion protein (Citrine is a YFP variant) (Miki et al., 2014) suggested that KCHa, b, and c, but not KHCd, are expressed in protonemal cells. However, since C-terminal tagging might perturb the function of the kinesin-14 subfamily, the motor domain of which is generally located closer to the C-terminus, we inserted the *Citrine* gene in front of endogenous *KCHa, b*, and *c* (elements other than the Citrine ORF were not integrated) (Figure S1A, B). With the newly selected Citrine-KCH lines, we confirmed that KCHa, b, and c are indeed expressed in protonemal cells, and furthermore, exhibit MT localisation at the cell tip (Figure 1C).

To test the contribution of KCH to nuclear positioning and other intracellular processes, we sequentially deleted four *KCH* genes by means of homologous recombination in the moss lines expressing GFP-tubulin and histoneH2B-mRFP (Figure S1C, D). The *KCHacd* triple knockout (KO) line grew in an indistinguishable manner to wild-type moss. However, when all four *KCH* genes were deleted, moss colony growth was severely retarded (Figure 1D, E). In addition, the gametophore leaf was curly and the rhizoid was much shorter than in the control line (Figure 1F). These phenotypes were suppressed when Cerulean-tagged KCHa was expressed by a constitutively active promoter in the quadruple KO line (Figure 1D–F). These results indicate that KCH is a critical motor in moss development, albeit not essential for moss viability.

### KCH is required for minus end-directed nuclear transport along MTs during interphase

We performed live imaging of the protonemal apical cells of the quadruple KO line. Unlike the *KCBP* KO line, sister nuclei moved towards the cell centre after chromosome segregation, indicating that minus end-directed motility during telophase was not impaired in the absence of KCH (Figure 2B, Movie 1). However, unlike the control cells that maintained cell-centre positioning of the nucleus during tip growth, the nucleus did not stop moving at the cell centre but migrated further towards the cell tip (apical cell) or moved back towards the cell plate (subapical cell). Consequently, the apical cells positioned the nucleus near the apical cell wall in the *KCH* KO line; this phenotype was rescued by ectopic Cerulean-KCHa expression (Figure 2C, D). During subsequent mitosis of apical cells, spindle assembly took place 10% more apically compared with the control line, resulting in an apical shift of the cell division site (p < 0.05, n = 13 [KO] and 7 [control]); this suggests a physiological role for nuclear positioning. Given the known MT polarity during interphase of apical cells (plus-ends predominantly face the apex; Hiwatashi et al., 2014, Yamada et al., 2017; Figure 2A) and the appearance of the nuclear migration defects contrary to kinesin-ARK depletion (the nucleus moves back to the cell plate in apical cells; Miki et al., 2015), it was suggested that KCH drives MT minus end-directed transport of the nucleus during interphase.

**Figure 2.**
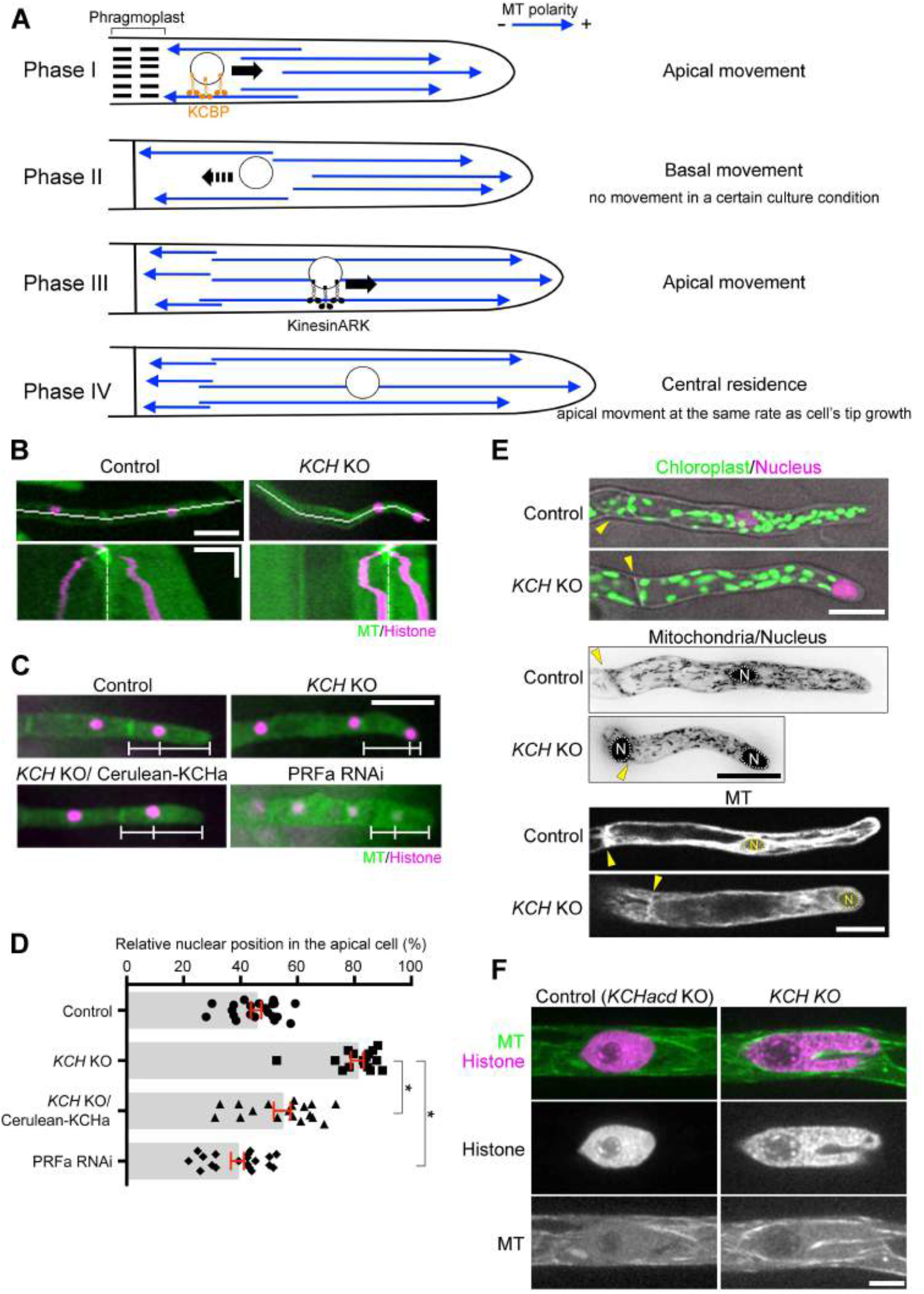
Nuclear positioning defects observed in KCH KO cells. **(A)** Schematic representation of MT polarity and nuclear positioning in apical cells based on previous studies (Hiwatashi et al., 2014; Miki et al., 2015; Yamada et al., 2017). **(B)** Nuclear migration defects were observed in the *KCH* KO line. Snapshot (top) and kymograph (bottom) of control and *KCH* KO cell are displayed. White lines in snapshots indicate the position where the kymograph was created. Kymograph starts from the mitotic metaphase. White broken lines indicate the position of the cell wall. Horizontal scale bar, 50 μm; vertical bar, 2 h. **(C)** Nuclei were apically localised in the absence of KCH, and the defect was rescued by Cerulean-KCHa expression. White bars indicate positions of the cell wall, nucleus, and apical tip. Scale bar, 50 μm. **(D)** Relative position of the nucleus within the apical cell was quantified. ‘0’ corresponds to the cell wall, whereas ‘100’ indicates the cell tip. Bars and error bars represent the mean and SEM, respectively. Control, n = 21, *KCH* KO, n = 16; *KCH* KO/Cerulean-KCHa, n = 18; PRFa RNAi, n = 19. *P < 0.0001 (unpaired t-test with equal SD, two-tailed). Note that induction of PRFa RNAi led to co-depletion of histoneH2B-mRFP, resulting in reduced histone signals (Nakaoka et al., 2012). Experiments and data analyses were performed twice, with data from one experiment displayed. **(E)** Distribution of nuclei, chloroplasts, mitochondria, and vacuoles (GFP-tubulin-excluded area). Yellow arrowhead and the “N” mark indicate the position of the cell wall and nucleus, respectively. Mitochondrial images were acquired with 5 z-sections (separated by 4 μm), and are displayed after maximum projection. Scale bars, 25 μm. **(F)** Nuclear deformation in the apical cell. Triple KO line (*KCHacd* KO) was used as control. Scale bar, 5 μm.

To determine whether the distribution of other organelles is perturbed in the absence of KCH, we assessed the intracellular distribution of chloroplasts (by autofluorescence), mitochondria (the N-terminal 78 amino acids of the γ-subunit of the *Arabidopsis* mitochondrial F_1_F_o_ ATPase) (Uchida et al., 2011; Nakaoka et al., 2015), and vacuoles (the GFP-tubulin-excluded areas). Unlike the nucleus, we did not observe any abnormal distribution or morphology of these organelles, suggesting that the effect of KCH deletion is specific to the nucleus (Figure 2E, Figure S2).

In *Arabidopsis* root and mesophyll cells, the nucleus is transported along actin filaments by the myosin XI-i motor, where its mutant exhibited not only nuclear dynamics defects but also nuclear deformation (the nucleus becomes rounder) (Tamura et al., 2013). Interestingly, when we observed the *KCH* KO line with spinning-disc confocal microscopy, we detected stretching and invagination of the nucleus in 55% (n = 29) of apical cells (Figure 2F). The discrepancy in nuclear shape in the *Arabidopsis myoXI-i* mutant and moss *KCH* KO is possibly because of the additional force applied on the nuclear surface of moss, for example, by kinesin-ARK. Nevertheless, this observation further supports the idea that KCH is responsible for nuclear motility.

### Processive motility of KCH *in vivo*

Cytoskeletal motor proteins that drive cargo transport are generally processive, where a single dimeric motor that attaches to a cytoskeletal filament (MT or actin) takes multiple steps towards one direction before dissociation. Some non-processive motors can also be transporters when multiple dimers participate in cargo transport. KCBP represents the latter example; multiple KCBP molecules bind to MTs via the motor domain and to vesicular cargo via the tail domain and execute long-distance transport (Yamada et al., 2017). The purified OsKCH1 motor was also shown to be non-processive but its cohort action drives actin motility along MTs (Walter et al., 2015). On the other hand, there have been contradictory reports as to whether full-length KCH shows processive motility *in vivo*. When tobacco GFP-NtKCH was expressed in tobacco BY-2 cells, motility of GFP signals (i.e. clustered GFP signals) along MTs was detected (Klotz and Nick, 2012). In contrast, GFP-AtKinG was observed only as static punctae on MTs in *Nicotiana benthamiana* leaf epidermal cells (Spiegelman et al., 2018). In these studies, however, GFP-tagged constructs were ectopically overexpressed and might not represent native KCH dynamics.

To test if endogenous *P. patens* KCH exhibits processive motility in moss cells, we acquired time-lapse images of Citrine-KCHa, which was expressed by the native promoter at the endogenous locus, using oblique illumination fluorescence microscopy. This microscopy allows for visualisation of the cortex-proximal region, which is Largely devoid of auto-fluorescence derived from chloroplasts. We observed punctate Citrine signals on MTs, and interestingly, the signals moved towards minus ends at a velocity of 441 ± 226 nm/s (± SD, n = 90) for 1.6 ± 1.5 μm (± SD, n = 74; Figure 3, Movie 2); this velocity was 8-folds faster than previously reported for NtKCH (Klotz and Nick, 2012). Since the microscopy technique employed is not sensitive enough to detect individual Citrine molecules (Jonsson et al., 2015), the diffraction-limited spots represent clustered Citrine-KCHa. This observation is consistent with the notion that KCH functions as a minus end-directed transporter of the nucleus.

**Figure 3.**
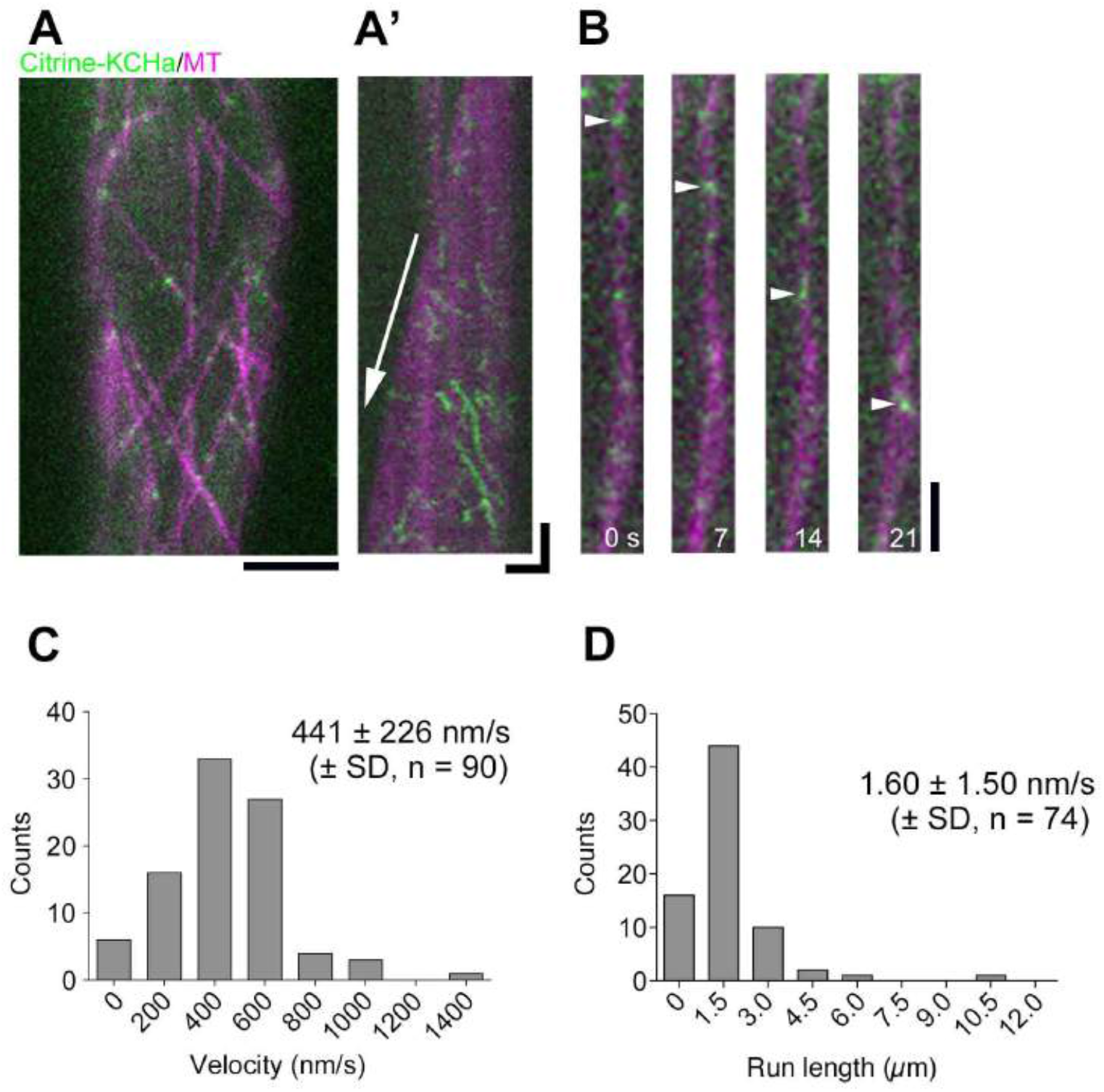
Processive, minus end-directed movement of KCH clusters along MTs in protonemal cells. **(A)** Localisation of Citrine-KCHa on endoplasmic MTs. Scale bar, 5 μm. **(A’)** Kymograph showing Citrine-KCHa signals moving towards the MT minus-end. Arrow indicates MT plus-end growth. Horizontal scale bar, 2 μm; vertical bar, 5 s. **(B)** Processive movement of Citrine-KCHa signals (arrowheads) along an endoplasmic MT. Scale bar, 2 μm. **(C, D)** Velocity and run length of moving Citrine-KCHa signals. Note that mean run length might be somewhat underestimated as some signals are photo-bleached during image acquisition.

### KCH promotes polarised tip growth

In addition to defects in nuclear transport, we observed severe tip growth retardation in the complete *KCH* KO line (Figure 4A, Movie 3). Moreover, abnormally branched tip cells were occasionally observed in the KO line, reflecting improperly polarised tip growth (Figure 4B). These defects were suppressed by Cerulean-KCHa expression, confirming that tip growth defects in the KO line were due to the loss of KCH proteins.

**Figure 4.**
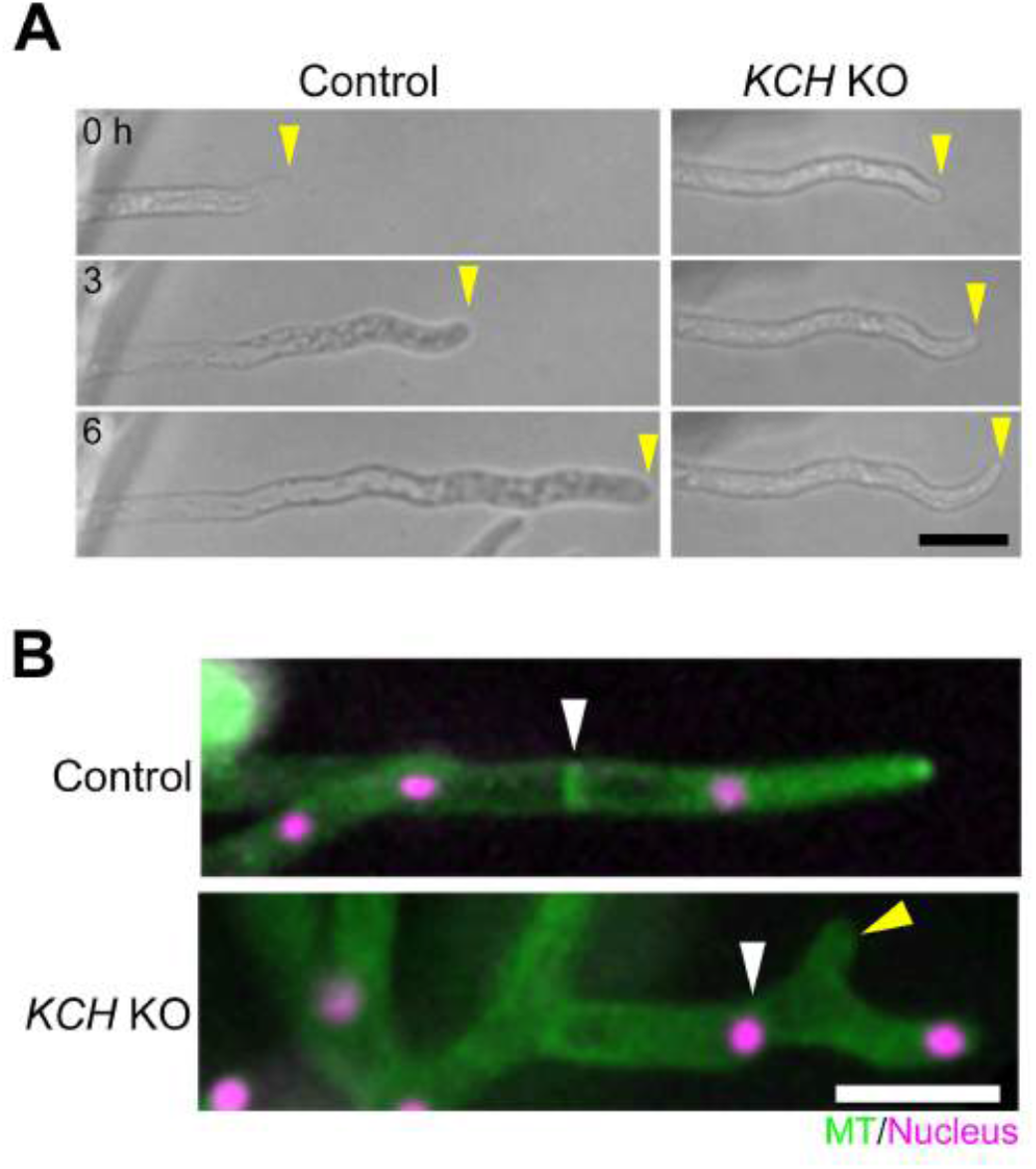
Protonemal tip growth is suppressed in *KCH* KO lines. **(A)** Tip growth was retarded in the absence of KCH. Yellow arrowheads indicate apical cell tips. **(B)** Branched cell observed in the *KCH* KO line. White and yellow arrowheads indicate the cell wall and abnormally branched tip, respectively. Scale bars, 50 μm.

The slow tip growth phenotype raised a possibility that nuclear mislocalisation may be a secondary effect of growth retardation in the *KCH* KO line. However, this was unlikely as the nucleus was mispositioned in the subapical cell, which hardly grows even in wild-type cells (Figure 2B, C). Nevertheless, to exclude this possibility, we examined nuclear positioning under another slow-growth condition generated by PRFa (profilin; an actin regulator) RNAi (Vidali et al., 2007; Nakaoka et al., 2012). We confirmed that the nucleus was more centrally localised when tip growth was suppressed following RNAi of PRFa (Figure 2C, D), suggesting that nuclear mistranslocation was not a secondary effect of the growth defect associated with *KCH* KO.

### MT focus formation at the tip requires actin and KCH

There are several tip-growing cells in plants, such as moss protonemata, root hairs, and pollen tubes. Actin filaments are essential for tip growth in these cells (Rounds and Bezanilla, 2013), whereas direction of growth is defined by MTs in some cases. In moss protonemata, MT disruption by inhibitors or depletion of key MT regulators leads to skewed or branched tip growth (Doonan et al., 1988; Hiwatashi et al., 2014). Since KCH binds to both MT and actin *in vitro* (Frey et al., 2009; Xu et al., 2009; Umezu et al., 2011; Walter et al., 2015; Tseng et al., 2018), we hypothesised that *P. patens* KCH might regulate cytoskeletal organisation at the cell tip.

We first performed live imaging of GFP-tubulin and lifeact-mCherry (actin marker) in control apical cells. As reported previously, MT and actin focal points were detected at the tip (Vidali et al., 2009; Hiwatashi et al., 2014); we also observed that they were largely—though not completely—colocalised with each other (Figure 5A, Movie 4). These focal points were not maintained when either MT or actin was disrupted by specific drugs (Figure 5A, Movie 4); thus, MT and actin focal points at the tip are mutually dependent. Interestingly, in the *KCH* KO line, the MT focus was smaller and much less persistent (Figure 5B, C, Movie 5). Furthermore, actin still accumulated at the transiently-formed MT focal point, indicating the presence of other factor(s) that crosslink MTs and actin at the tip (Figure 5D).

**Figure 5.**
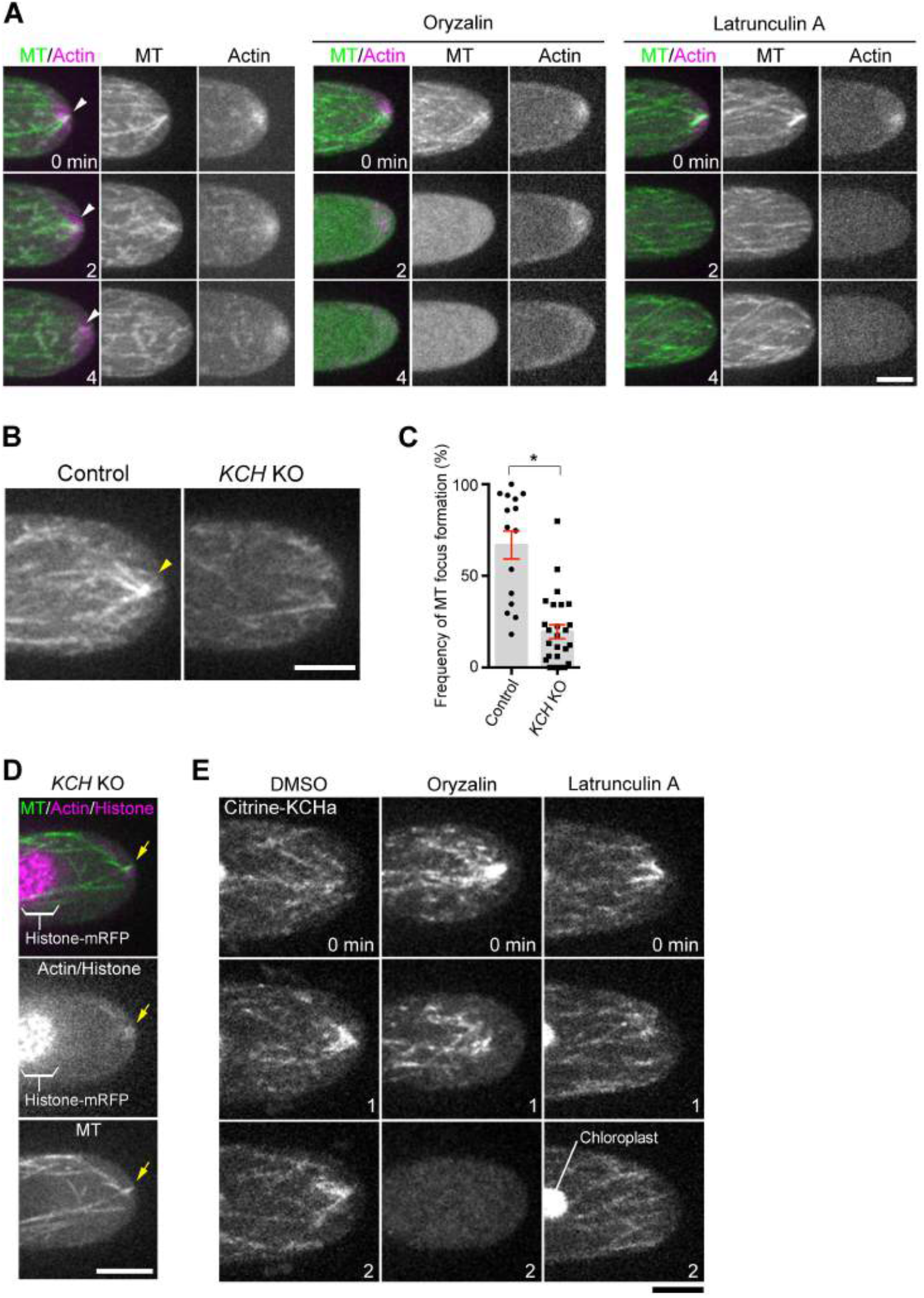
MT focusing at the apical cell tip requires actin filaments and KCH. **(A)** Formation of MT and actin foci (arrowheads) at the tip is mutually dependent. The tip of a caulonemal cell expressing GFP-tubulin (MT) and lifeact-mCherry (actin) was observed after oryzalin or latrunculin A treatment. Actin focal points were not maintained at the apical cell tip after oryzalin addition, whereas MT focal points were rarely observed after latrunculin A treatment. **(B)** Persistent formation of the MT focus (arrowheads) at the apical cell tip is dependent on KCH. **(C)** Frequency of MT focus formation. Images were acquired and analysed every 3 s for 5 min. Bars and error bars represent the mean and SEM, respectively. Control, n = 15; *KCH* KO, n = 26. *P < 0.0001 (unpaired t-test with equal SD, two-tailed). Observations were performed independently three times, and the data analysed twice. The combined data is presented. **(D)** Actin (lifeact-mCherry) accumulation around the transiently-formed MT focus (GFP-tubulin) in the absence of KCH. **(E)** Actin-independent localisation of Citrine-KCHa on MTs. Drugs were added at time 0. Note that signals derived from chloroplast auto-fluorescence are visible in some panels. Scale bars, 5 μm.

KCHa localisation at the tip disappeared following MT destabilisation by oryzalin treatment, whereas actin destabilisation by latrunculin A did not affect KCHa MT association (Figure 5E); thus, KCH and MT co-localise independent of actin, but focal points cannot be formed without actin.

### The C-terminal region of KCH is required for tip growth but not for nuclear transport, whereas the CH domain is dispensable for either activity

To functionally dissect the KCH protein, we constructed several truncation constructs tagged with Cerulean, transformed them into the *KCH* KO line, selected for transgenic lines, and assessed if the phenotypes were rescued (Figure 6A, S1E). Surprisingly, the truncated KCHa deleted of the canonical CH domain, which has been assumed to be the actin-binding site, restored protonemal colony growth and rhizoid development (Figure 6B, C). The gametophore leaves also showed considerable recovery of morphology (Figure 6B). In contrast, we could not obtain a transgenic line that showed normal protonemal growth or gametophore/rhizoid development after transformation of the KCH fragment with a deleted C-terminal extension (Figure 6B, C).

**Figure 6.**
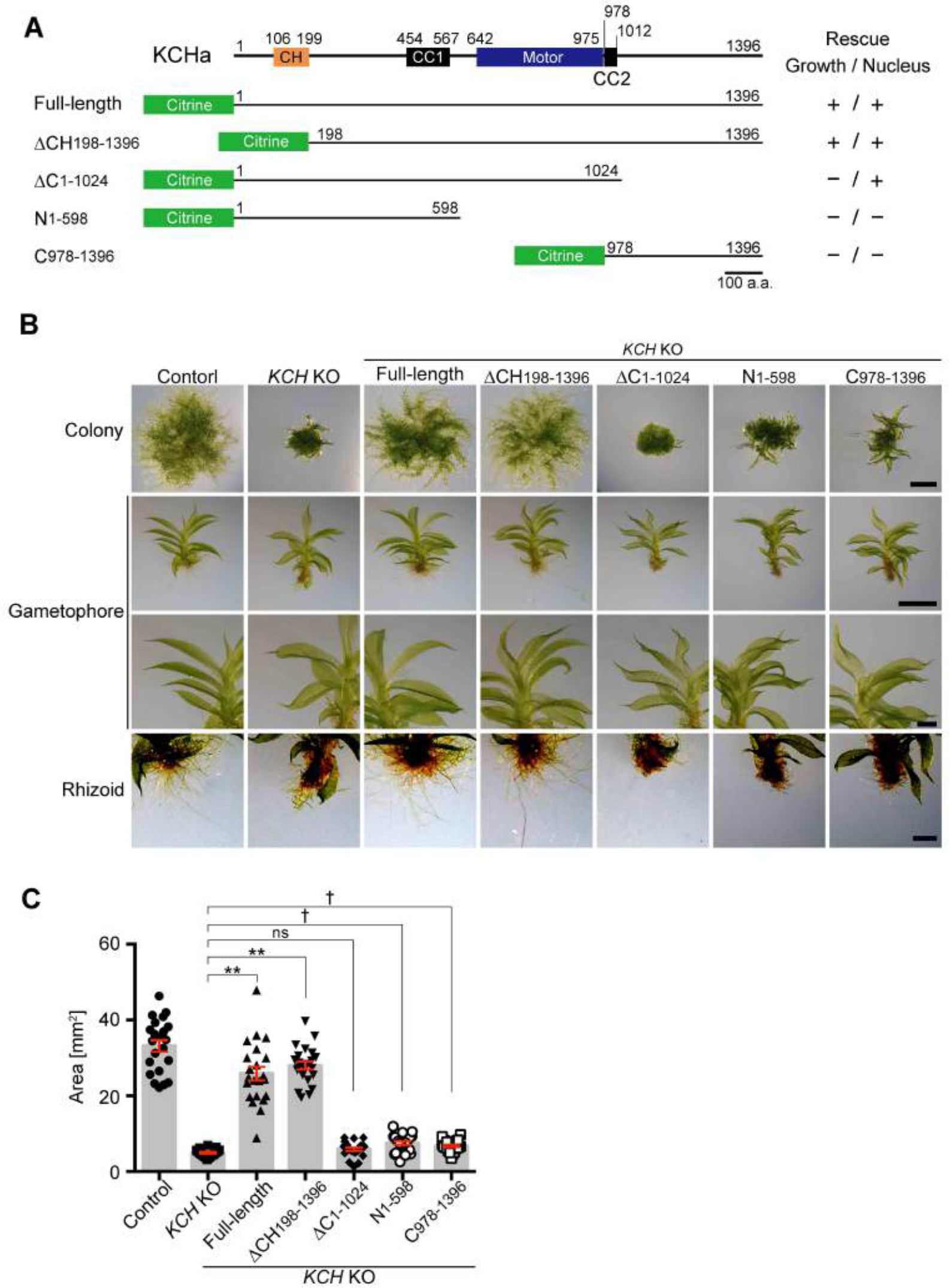
Mapping of KCH functional domains for moss development. **(A)** Schematic representation of four truncation constructs and their ability to rescue defects in nuclear transport or protonemal growth. Citrine was attached to the N-terminus of each fragment. **(B)** Protonemal growth, gametophore leaf morphology, and rhizoid development after expression of truncated constructs in the *KCH* KO line. Scale bars, 2 mm (top two rows) or 0.5 mm (bottom two rows). **(C)** Colony size comparison between control (parental line expressing GFP-tubulin/histoneH2B-mRFP), *KCH* KO, and *KCH* KO/Citrine-KCHa truncations. Bars and error bars represent the mean and SEM, respectively. Control, n = 22; *KCH* KO, n = 21; *KCH* KO/Citrine-FL (1-1396), n = 22; *KCH* KO/Citrine-ΔCH (198-1396), n = 22; *KCH* KO/Citrine-ΔC (1-1024), n = 22; *KCH* KO/Citrine-N (1-598), n = 22; *KCH* KO/Citrine-C (978-1396), n = 22. **P < 0.0001; ns (not significant), P > 0.06 (unpaired t-test with equal SD, two-tailed). fAlthough colony size was slightly but significantly larger than that of the KO line in this experiment, it was slightly smaller in another experiment, indicating that these two short fragments did not rescue colony growth. Experiments were performed three times and the data analysed twice. The data of one experiment is displayed.

At the cellular and intracellular levels in the protonemata, tip growth (Figure 7A, Movie 3), persistent MT focus formation (Movie 6), and nuclear positioning (Figure 7B, C) were restored by ∆CH expression. In contrast, we observed recovery of nuclear positioning, but not tip growth or MT focus formation, in the ∆C lines, indicating that the C-terminal extension is dispensable for nuclear transport function but is essential for tip growth.

**Figure 7.**
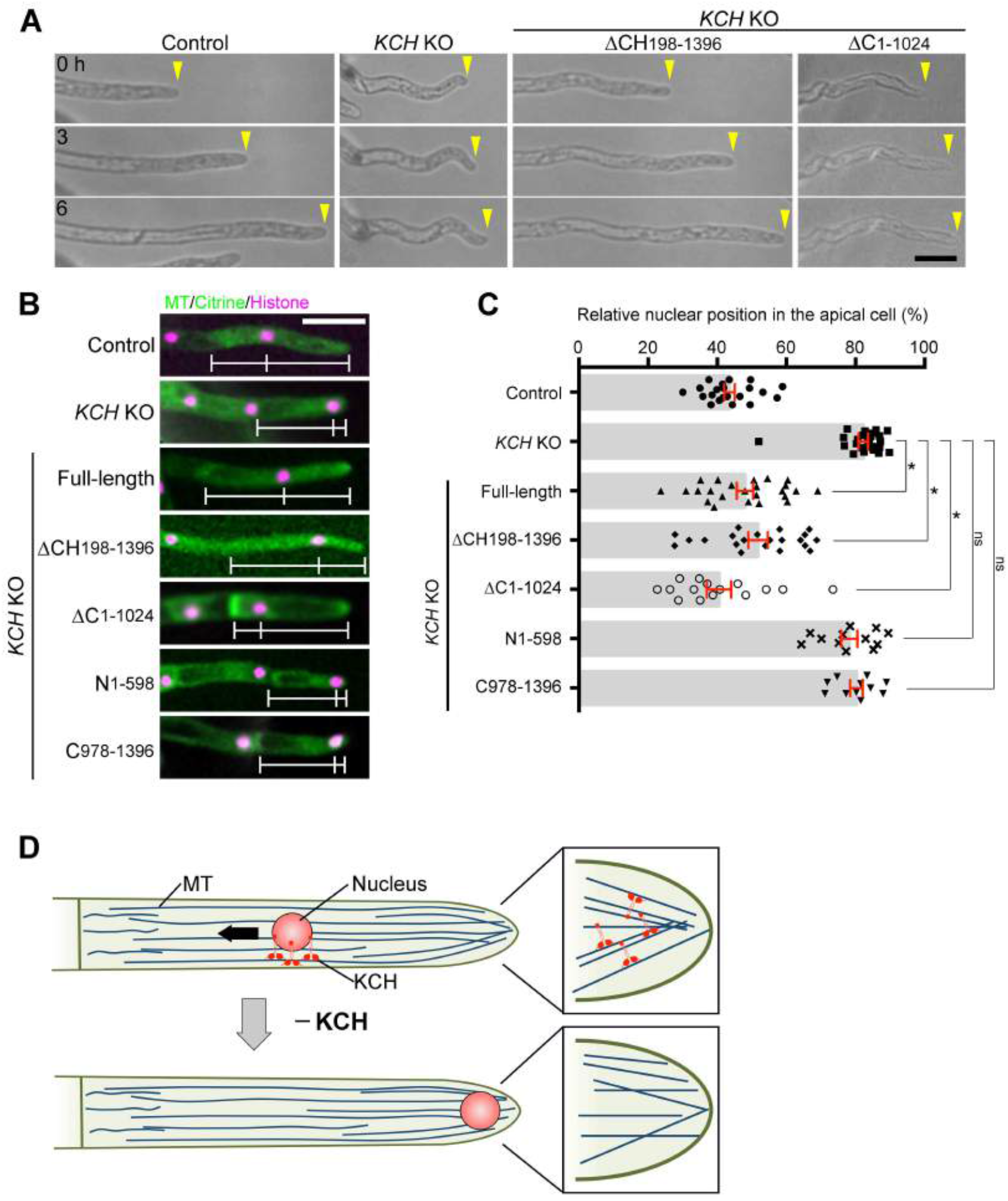
C-terminal extension of KCH is critical for tip growth, but not for nuclear transport. **(A)** Tip growth was assessed after expression of the truncated KCH in the KO line. Scale bar, 50 μm. **(B)** Nuclear positioning after truncated protein expression in the KCH KO line. White bars indicate the positions of the cell wall, nucleus, and apical tip. Scale bar, 50 μm. See Movie 6 for high-resolution MT images in some transgenic lines. **(C)** Relative position of the nucleus within the apical cell was quantified for each line. ‘0’ corresponds to the cell wall, whereas ‘100’ indicates the cell tip. Bars and error bars represent the mean and SEM, respectively. Control, n = 22; *KCH* KO, n = 26; *KCH* KO/Citrine-FL (1-1396), n = 24; *KCH* KO/Citrine-ΔCH (198-1396), n = 21; *KCH* KO/Citrine-ΔC (1-1024), n = 15; *KCH* KO/Citrine-N (1-598), n = 12; *KCH* KO/Citrine-C (978-1396), n = 11. *P < 0.0001; ns (not significant), P > 0.1 (unpaired t-test with equal SD, two-tailed). Observations were performed independently twice or more times, and the data analysed twice. The data of one experiment is displayed. **(D)** Role of KCH kinesin for tip cell growth and nuclear transport in P. patens. KCH is a minus-end-directed kinesin required for retrograde transport of the nucleus (arrow). KCH is also critical for tip growth as it stably focuses MTs at the apical tip. Actin is enriched around the MT focus (not displayed in this cartoon).

## DISCUSSION

We generated, for the first time, a plant completely lacking in KCH proteins, which exhibited several noticeable phenotypes. We focused our study on two prominent phenotypes associated with protonemal apical cells, nuclear mispositioning and tip growth retardation. This study also represents the first report, to our knowledge, on intracellular dynamics of function-verified KCH protein expressed from the native locus. Our data elucidated the function of KCH as a long-distance retrograde transporter and its role in cytoskeletal coalescence, during which an unanticipated molecular mechanism might be involved (Figure 7D).

### KCH is a potent retrograde transporter

The dimeric, truncated form of rice KCH1 or moss KCHa was shown to be non-processive in *in vitro* motility assays (Jonsson et al., 2015; Walter et al., 2015), whereas a recent report showed processive motility of rice KCH2 as a dimer; OsKCH2 possesses unique sequences adjacent to the C-terminus of the motor domain which ensure processivity (Tseng et al., 2018). However, a cohort action of rice KCH1 can transport actin filaments along MTs over long distances, suggesting that KCH family proteins can generally function as long-distance cargo transporters (Walter et al., 2015). Our observation of endogenous KCHa dynamics in the cytoplasm indicated that this kinesin forms a cluster and indeed moves processively towards the minus ends of MTs. Its run velocity (~440 nm/s) was approximately 3-folds faster than that of ATK (kinesin-14-I) and comparable to KCBP (kinesin-14-VI, 413 nm/s)—two other kinesin-14 family proteins for which processive motility in clusters and cargo transport function have been identified (Jonsson et al., 2015; Yamada et al., 2017). These results suggest that KCH proteins are potent minus end-directed transporters.

Paradoxically, a minus-end-directed motor is enriched at the MT plus end at the cell tip (Figure 1C, 5E). This might be achieved by interacting with other MT plus end-tracking proteins or actins. However, the results obtained using oblique illumination fluorescence microscopy indicate that many KCH molecules associate only transiently with the MT and then dissociate before moving along the MT (Movie 2). Therefore, it is possible that both non-motile and motile proteins can be visualized using confocal microscopy at the MT-rich cell tip. Consistent with this idea, the mCherry-tubulin signals increased at the tip similar to Citrine-KCH (Figure 1C), and the latter localised uniformly to the MT when the MT focus was disrupted by latrunculin A treatment (Figure 5E, right). Whether minus end-directed motility of KCH is required for cytoskeletal organisation at the tip remains to be determined.

### KCH drives nuclear transport

Our study identified a defect in nuclear positioning in the absence of KCH. Nuclear translocation during the cell cycle of protonemal apical cells can be divided into four phases (Figure 2A), each driven by MT-based transport. KCBP is a minus end-directed kinesin required for nuclear transport specifically during phase I, the post-mitotic phase (Yamada et al., 2017). In contrast, kinesin-ARK is responsible for plus end-directed transport during phase III and possibly also phase IV, which correspond to the majority of the interphase (Miki et al., 2015). KCH is the third kinesin identified required for nuclear positioning; its KO phenotype indicated that KCH also functions during IV. The following three pieces of data strongly suggest that KCH actually transports the nucleus, rather than indirectly affecting nuclear positioning by, for example, altering overall MT polarity; (1) KCH showed minus end-directed, processive motility in cells (Figure 3), (2) the nucleus moved all the way to the cell tip in the *KCH* KO line (Figure 2B), where plus ends of MTs are enriched regardless of KCH presence (Figure 5B), and (3) in the absence of KCH, abnormal localisation was detected for the nucleus, but not for the other three cytoplasmic organelles (Figure 2E). Thus, we envisage that kinesin-ARK and KCH execute bi-directional transport of the nucleus, reminiscent of animal cells where antagonistic cytoplasmic dynein and kinesin-1 (or kinesin-3) motors are involved (Tanenbaum et al., 2010; Tsai et al., 2010). Consistent with this model, the nucleus unidirectionally moved towards the cell plate or cell tip in the absence of kinesin-ARK or KCH, respectively. We could not detect enrichment of KCH or kinesin-ARK (Miki et al., 2015) on the nuclear surface during microscopy; however, failure in detection does not necessarily preclude the possibility that KCH acts as a transporter, as the fluorescence signals might not be distinguishable from cargo-free motors in the cytoplasm or cytoplasmic background signals. This is particularly a likely scenario in moss protonemata, since auto-fluorescence derived from chloroplasts is dominant in the cytoplasm. Using oblique illumination microscopy, we observed Citrine-KCHa signals for each cortex-proximal MT, suggesting that KCH also associates with nucleus-proximal MTs (note that the nucleus cannot be located in the focal plane using this microscopy technique). Our truncation/rescue experiments suggest that the region downstream of the CH domain and upstream of the motor domain is required for nuclear attachment. It would be interesting to search for the nuclear envelope-associated adaptor of KCH in future studies.

### KCH and actin ensure MT focus formation for tip growth

Apical tip growth was severely suppressed in the *KCH* KO line. Based on previous studies, this phenotype could be attributed to defects in the actin cytoskeleton, MT cytoskeleton, and/or lipid biogenesis at the tip (van Gisbergen et al., 2012; Vidali and Bezanilla, 2012; Hiwatashi et al., 2014; Saavedra et al., 2015). Defects in general housekeeping processes such as protein translation would also perturb cell growth. Although involvement of KCH in lipid production or other general cellular processes was not excluded, our localisation and phenotypic data more strongly suggest that KCH regulates MT and possibly also actin cytoskeletons for tip growth. Most compellingly, the characteristic focus composed of MTs—which we observed to largely coincide with the actin focal point—did not persistently form in *KCH* KO cells. However, unlike for actin or MT destabilisation, tip growth was not completely inhibited in KO cells. Phenotypically, this was consistent with the observation that smaller and transient MT/actin foci were still detectable in the KO line. Residual MT bundling activity may be mediated by other proteins such as plant-specific plus end-directed kinesins KINIDa and b, whose double deletion resulted in curved growth accompanied with MT focus impersistence (Hiwatashi et al., 2014). In addition, the mislocalised nucleus also dampened tip growth, as this large organelle occasionally occupied the apical space where MT and actin normally form clusters (e.g. Figure 2B). Thus, another significant role of nuclear positioning in plants—other than division site determination—may be ensuring proper organisation of the cytoskeletal network for cell function. Nevertheless, nuclear mislocalisation is not the sole factor contributing to tip growth suppression, since MT foci were not stably maintained in KO cells whose nuclei resided fairly distant from the tip (e.g. Figure 5B). Furthermore, the C-terminal deletion construct restored nuclear positioning but not growth defects. Thus, cytoskeletal disorganisation independent of nuclear positioning is likely the major factor responsible for growth retardation in *KCH* KO cells.

The mechanism via which KCH promotes the formation of a large and persistent ensemble of the two cytoskeletal filaments remains unclear. MT-actin coalescence in the *KCH* KO line indicates the presence of other protein(s) that bridge the two filaments. KCH might support the coalescence via MT crosslinking. Alternatively, the model that KCH, independent of the CH domain, coalesces MTs with actin by directly binding to both filaments remains viable; intriguingly, an *in vitro* study indicates that actin interacts also with the motor domain of rice KCH-O12 (Umezu et al., 2011). Furthermore, the C-terminal extension of KCH was critical for MT coalescence. Long (> 150 a.a.) C-terminal extension from the motor domain is unique to the plant kinesin14-II– V families, and apart from the coiled-coil, no sequences have been identified in this region from which protein function can be deduced. Our study showed that this unusual extension is a critical element of KCH, yet its exact function remains unclear; it may be required for KCH accumulation at the tip or it may constitute an additional MT or actin interaction interface.

### Conservation of KCH function

In flowering plants, KCH family proteins show more divergence in amino acid sequences than in moss (Figure 1A), and each member appears to play a distinct role. Interestingly, several reported phenotypes associated with KCH depletion and overexpression in flowering plants appear to be consistent with our findings for moss. For example, coleoptile cells were shorter in the rice *kch1* mutant, whereas tobacco BY-2 cells were elongated upon overexpression of rice KCH (Frey et al., 2010). Tobacco GFP-KCH, when ectopically expressed, decorates cortical MT arrays but is also abundantly present around the nucleus (Klotz and Nick, 2012). Furthermore, overexpression of rice KCH in the tobacco BY-2 cell line induced nuclear positioning defects (Frey et al., 2010). Although these studies did not elucidate the basis of the phenotypes, our study suggests that KCH may transport the nucleus and promote MT-actin interaction in flowering plants as well. However, KCH functions might not be limited to nuclear transport and MT-actin interactions during moss development, as many KCH proteins not associated with the nucleus were observed running along MTs (Figure 3). The complete KO line generated in this study is a potentially valuable resource for uncovering the list of KCH functions throughout plant development.

## METHODS

### Moss culture and transformation

Moss lines used in this study are listed in Table S1; all lines originated from the *Physcomitrella patens* Gransden2004 strain. GFP-tubulin/histoneH2B-mRFP and mCherry-tubulin strains were used as mother strains for gene disruption and Citrine tagging, respectively (Nakaoka et al., 2012; Kosetsu et al., 2013). Methodologies of moss culture, transformation, and transgenic line selection were previously described thoroughly (Yamada et al., 2016). Briefly, cells were cultured on BCD agar medium for imaging. Transformation was performed by the standard PEG-mediated method and stable lines were selected with antibiotics (blasticidin S, nourseothricin, and hygromycin). Gene knockouts were obtained by replacing endogenous *KCH* genes with a drug-resistant marker flanked by lox-P sequences. After deleting *KCH-a* and *b*, the two markers were removed by transient expression of Cre recombinase, followed by replacement of *KCH-d* and *b* with the same markers. The *Citrine* gene was inserted into the N-terminus of *KCH-a* via homologous recombination. Drug-resistant genes were not integrated into the genome, as non-linearized, unstable plasmids containing a drug resistant marker were co-transformed with the linearized plasmid containing Citrine tag constructs for transient drug selection. Gene disruption and Citrine tag insertion were confirmed by PCR.

### Plasmid construction

Plasmids (and primers for plasmid construction) used for gene disruption, protein expression, and Citrine tagging are listed in Table S2. Gene knockout constructs were designed to replace endogenous *KCH* genes with a drug-resistant marker flanked by lox-P sequences. One kb of genomic DNA sequences upstream/downstream of start/stop codons were amplified and cloned into the vectors containing lox-P sequences and drug-resistant markers. To generate a plasmid for Citrine tagging, 1 kb of genomic DNA sequences upstream/downstream of the *KCH* gene start codon were cloned into the pKK138 vector (Kosetsu et al., 2013; Yamada et al., 2016). To generate truncation/rescue plasmids, *KCH-a* sequences were amplified from a cDNA library and ligated into the pENTR/D-TOPO vector containing *Cerulean* or *Citrine* sequences, followed by a Gateway LR reaction into a pTM153 vector that contains the EF1α promoter, blasticidin-resistance cassette, and PTA1 sequences designated for homologous recombination-based integration (Miki et al., 2016). Lifeact-mCherry was expressed by the actin promoter.

### In vivo microscopy

Methods for epifluorescence and spinning-disc confocal microscopy were previously described thoroughly (Yamada et al., 2016). Briefly, protonemal cells were plated onto glass-bottom plates coated with BCD agar medium and cultured for 4–5 d. The KO line was pre-cultured on BCD medium covered with cellophane for 2– 3 d before being placed onto glass-bottom plates. Long-term imaging by a wide-field microscope (low magnification lens) was performed with a Nikon Ti (10× 0.45 NA lens and EMCCD camera Evolve [Roper]). High-resolution imaging was performed with a spinning-disc confocal microscope (Nikon TE2000 or Ti; 100× 1.45 NA lens, CSU-X1 [Yokogawa], and EMCCD camera ImagEM [Hamamatsu]). Oblique illumination microscopy was performed as previously described (Jonsson et al., 2015; Nakaoka et al., 2015); cells were cultured in BCDAT medium covered with cellophane and imaging was performed with a Nikon Ti microscope with a TIRF unit, a 100× 1.49 NA lens, GEMINI split view (Hamamatsu), and EMCCD camera Evolve (Roper). All imaging was performed at 24–25°C in the dark, except for the protonema growth assay that utilises light.

### Drug assay

Mosses were plated on agar gel in 35-mm dishes following standard procedure and incubated for 4–5 d (Yamada et al., 2016). Prior to drug treatment, gels were excised from the dish except the central ~1 cm^2^ area on which moss colonies grew. Then, 1.5 mL water was added to the dish and incubated for 30 min (i.e. equilibrated), followed by addition of 1.5 mL drug solution. Cells were treated with 10 μM oryzalin (AccuStandard), 25 μM latrunculin A (Wako) or 0.5–1% DMSO as control.

### Colony growth assay

Protoplasts prepared by the standard driselase treatment were washed three times with 8% mannitol solution, followed by overnight incubation in protoplast liquid medium in the dark (Yamada et al., 2016). Protoplast solution was mixed with PRM-T and plated onto a cellophane-covered PRM plate (cellophane was a gift from Futamura Chemical, Co., LTD.) and incubated for 4 d. The cellophane was transferred to a fresh BCDAT plate, incubated for 5 d, and then colonies were picked and used to inoculate a yet new BCDAT plate. After incubation for a further 10–11 d, colonies were imaged with a commercially-available digital camera (Olympus C-765) or stereoscopic microscope (Nikon SMZ800N and Sony ILCE-QX1α).

### Phylogenetic tree construction

Following the method described in (Miki et al., 2014; Miki et al., 2015), kinesin sequences were aligned with MAFFT and the gaps were removed from the alignment using MacClade. The phylogenetic tree was constructed using neighbour joining (NJ) methods using the Molecular Evolutionary Genetics Analysis (MEGA) software. Statistical support for internal branches by bootstrap analyses was calculated using 1,000 replications.

### Data analysis

To quantify the relative position of the nucleus in the apical cell, microscope images that showed the cell wall, nucleus, and cell tip were analysed manually with ImageJ. Non-mitotic cells were randomly selected. The velocity and run length of Citrine-KCHa motility along endoplasmic MTs were quantified based on kymographs, which were generated from the images acquired with oblique illumination fluorescence microscopy. To measure the area of moss colonies, cultured moss colony images were acquired by a digital camera (Olympus C-765). All acquired images were aligned side-by-side and the generated single image was analysed with ImageJ; specifically, the image was processed by ‘Make binary’ and the colonies in the processed image were automatically detected and measured by ‘Analyze particles’ (size, 1.0-Infinity [mm^2^]; circularity, 0.00-1.00). To quantify the duration of MT focus formation at the tip, GFP-tubulin was imaged every 3 s and the presence or absence of MT foci was manually judged at each time frame; the focus was recognised when two or more MT ends were converged.

### Accession numbers

Sequence data used in this article can be found in the Phytozome database under the following accession numbers, *KCHa*, Pp3c14_19550; *KCHb*, Pp3c2_9150; *KCHc*, Pp3c17_21780; *KCHd*, Pp3c24_19380.

## ACKNOWLEDGMENTS

We thank Mitsuyasu Hasebe for providing the plasmids, Momoko Nishina, Tomohiro Miki, and Rie Inaba for technical assistance, as well as Tomomi Kiyomitsu, Elena Kozgunova, Peishan Yi, and Tomohiro Miki for helpful comments regarding the manuscript. This work was funded by the TORAY Science Foundation and JSPS KAKENHI 15K14540 and 17H06471 (to G.G). M.Y. is a recipient of a JSPS pre-doctoral fellowship.

## AUTHOR CONTRIBUTIONS

M.Y. and G.G. conceived and designed the research project. M.Y. performed the experiments and analysed the data. M.Y. and G.G. wrote the paper.

## MOVIE LEGENDS

**Movie 1.** Nuclear migration defects in the KCH KO line

Images were acquired every 3 min with epifluorescence microscopy. Green, MT; magenta, histone. Scale bar, 50 μm.

**Movie 2. Processive motility of clustered Citrine-KCH-a**

Citrine (green) and mCherry-tubulin (magenta) images were acquired every 0.24 s in the endoplasm of a protonemal cell. Note that auto-fluorescence derived from chloroplasts are also visible (green). Scale bar, 5 μm.

**Movie 3. Tip growth retardation in the KCH KO line**

Images were acquired every 3 min (10× lens, transmission light). Expression of the CH-deleted construct restored normal growth, whereas C-terminal deletion did not. Scale bar, 50 μm.

**Movie 4. MT and actin focal points at the cell tip**

Images were acquired every 30 s with spinning-disc confocal microscopy. Drugs were added at time 0. Green, MT; magenta, actin. Scale bar, 5 μm.

**Movie 5. KCH-dependent formation of the MT focus at the apical cell tip**

GFP-tubulin images were acquired every 3 s with spinning-disc confocal microscopy. Scale bar, 5 μm.

**Movie 6. MT focus at the apical cell tip after expression of truncated KCH**

GFP-tubulin images were acquired every 3 s with spinning-disc confocal microscopy. Note that signals derived from Citrine-KCH fragments are also merged in the rescue samples, since the emission filter associated with our microscope could not separate Citrine fluorescence from GFP. Scale bar, 5 μm.

---

**Figure S1.**
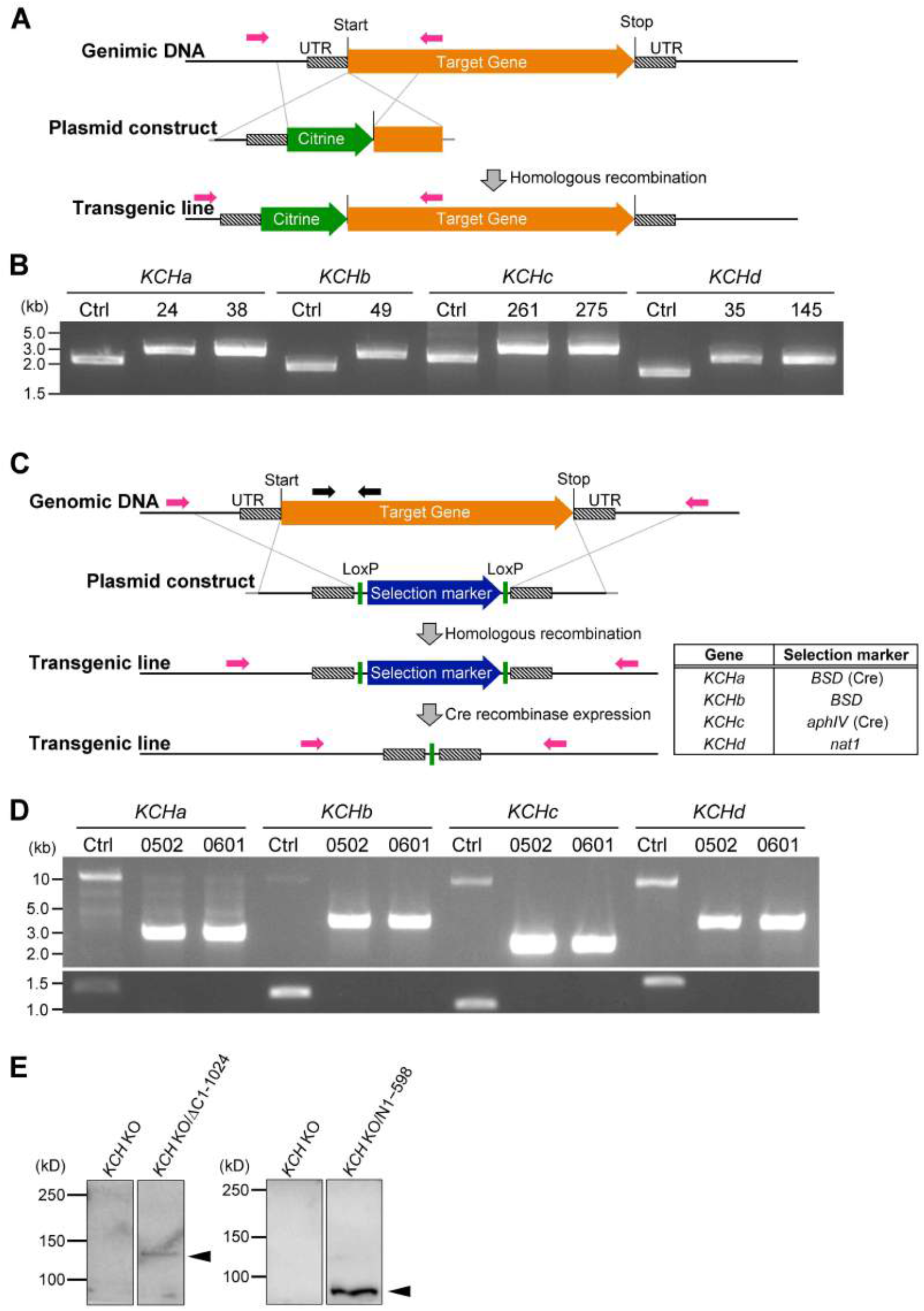
*KCH* gene tagging and disruption via homologous recombination. **(A, B)** Confirmation of Citrine-KCH lines by genotyping PCR. Genomic DNA from the control line expressing and Citrine-KCH lines (clone numbers are indicated at the top of the gel) were used as PCR templates.Citrine-KCH lines were generated from the control line expressing mCherry-tubulin. PCR primers are indicated as magenta arrows. No selection markers were integrated. **(C, D)** Confirmation of the quadruple *KCH* KO line by genotyping PCR. Genomic DNA from the control line and two *KCH* KO lines (#0502 and #0601) were used as PCR templates. *KCH* KO was generated from the control (‘Ctrl’) line expressing GFP-tubulin and histoneH2B-mRFP. Top and bottom panels show ‘Outer’ PCR and ‘Inner’ PCR, respectively. ‘Outer’ PCR primers (magenta arrows) were designed to flank the homologous recombination arm of each gene, whereas ‘Inner’ primers (black arrows) were designed for the open reading frame. Selection markers used are indicated in the table on the right. **(E)** Confirmation of Citrine-tagged truncated protein expression by immunoblotting with the anti-GFP antibody. Other fragments, including full-length Citrine-KCH, could not be detected as specific bands above the background signals.

**Figure S2.**
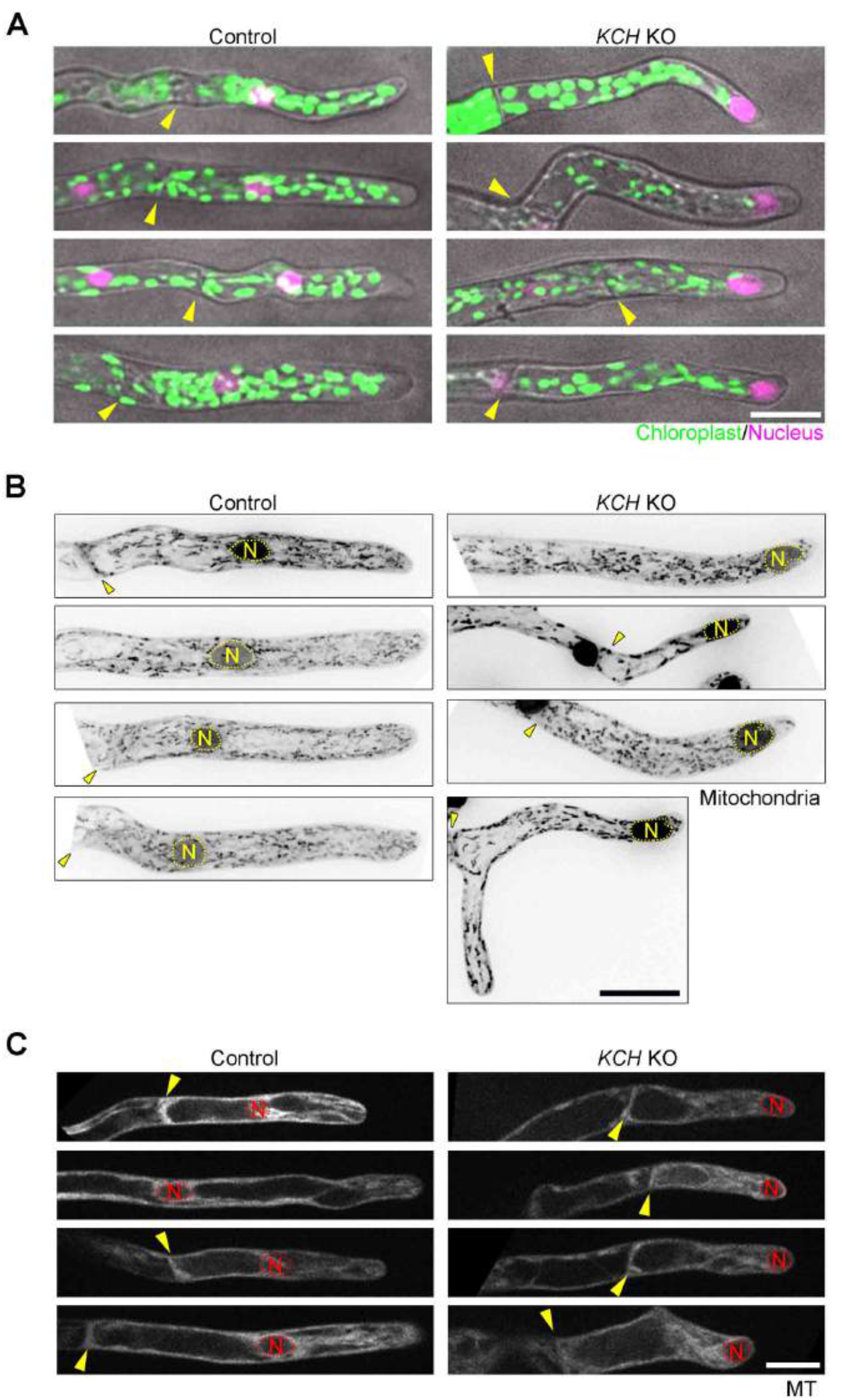
Organelle distribution in the *KCH* KO line. **(A-C)** Distribution of nuclei, chloroplasts, mitochondria, and vacuoles (GFP-tubulin-excluded area). Four cells were randomly picked and displayed for each organelle. Yellow arrowhead and the “N” mark indicate the position of the cell wall and nucleus, respectively. Mitochondrial images were acquired with 5 z-sections (separated by 4 μm), and are displayed after maximum projection. Scale bars, 25 μm.

**Table S1.**
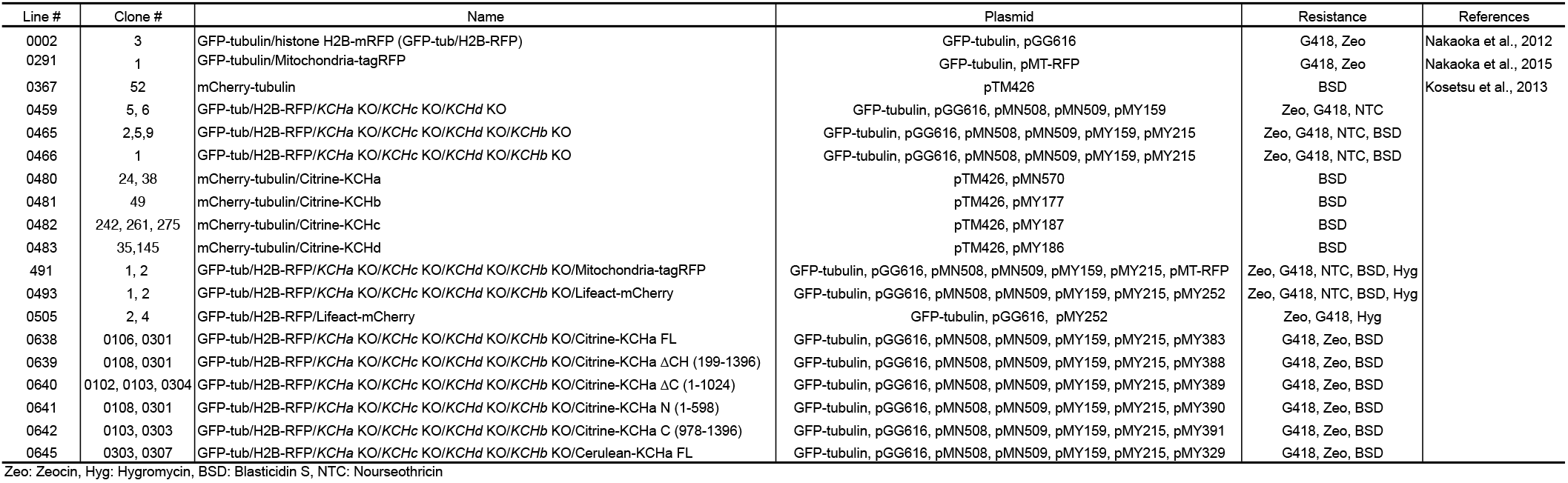
Moss lines used in this study

**Table S2.**
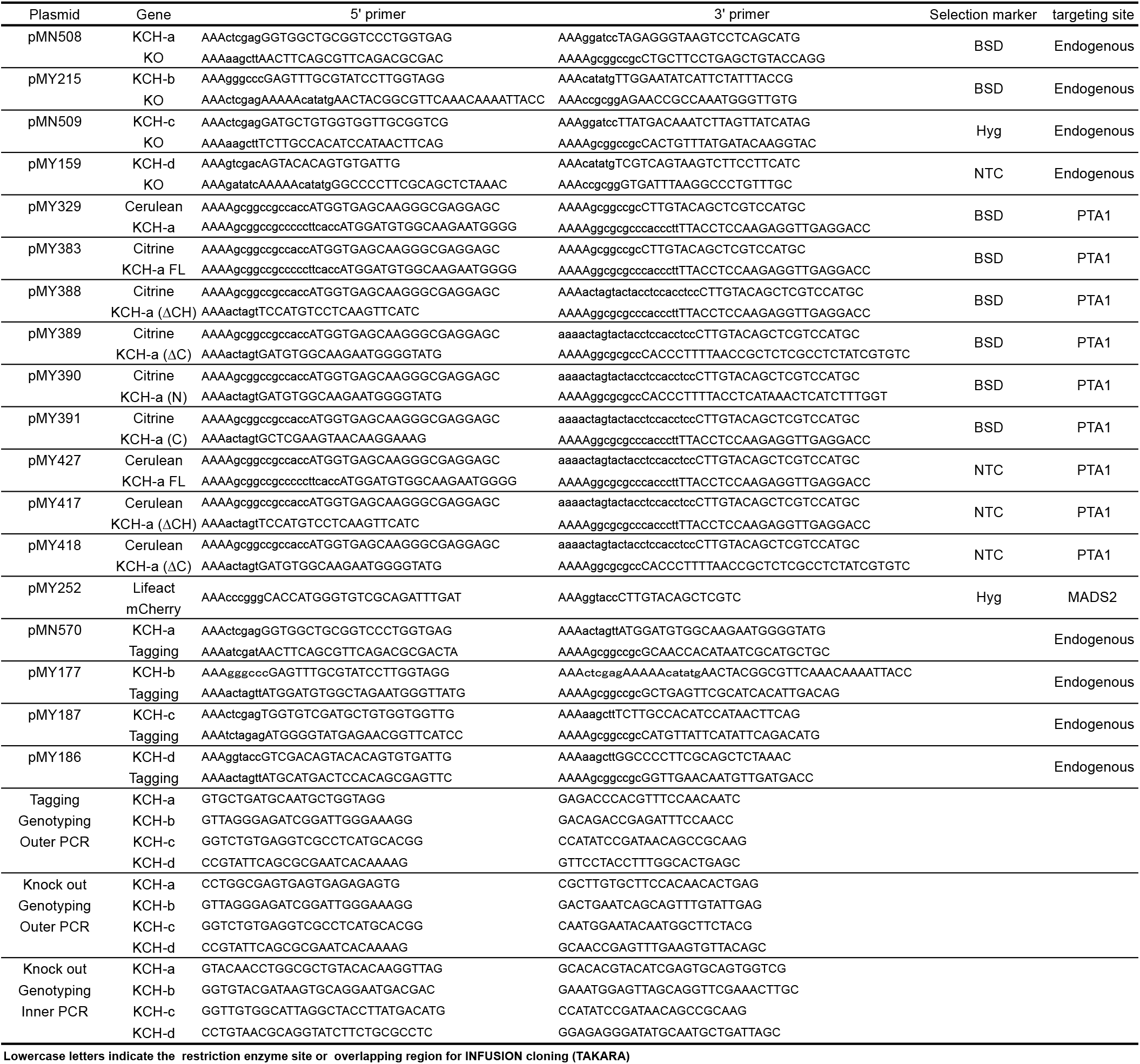
List of PCR primers and plasmids used in this study

